# mRNA with mammalian codon bias accumulates in yeast mutants with constitutive stress granules

**DOI:** 10.1101/2019.12.19.881961

**Authors:** Natalia V. Kozlova, Chantal Pichon, A. Rachid Rahmouni

**Affiliations:** Centre de Biophysique Moléculaire, UPR 4301 du CNRS, Rue Charles Sadron, 45071 Orléans, France; Université d’Orléans, Colléguim Sciences et Techniques, 45071 Orléans, France

**Keywords:** stress granules, *Saccharomyces cerevisiae*, mRNA accumulation

## Abstract

Stress granules and P bodies are cytoplasmic structures assembled in response to various stress factors and represent sites of temporary storage or decay of mRNAs. Depending on the source of stress, the formation of these structures may be driven by distinct mechanisms, but several stresses were shown to stabilize mRNAs via inhibition of deadenylation. A recent study identified yeast gene deletion mutants with constitutive stress granules and elevated P bodies; however, the mechanisms which trigger its formation remain poorly understood. Here, we investigate the possibility of accumulating mRNA with mammalian codon bias, which we termed the model RNA, in these mutants. We found that the model RNA accumulates in *dcp2* and *xrn1* mutants and in four mutants with constitutive stress granules overlapping with P bodies. However, in eight other mutants with constitutive stress granules the model RNA is downregulated, or its steady state levels vary. We further suggest that the accumulation of the model RNA is linked to its protection from the main mRNA surveillance path. However, there is no obvious targeting of the model RNA to stress granules or P bodies. Thus, accumulation of the model RNA and formation of constitutive stress granules occur independently and only some paths inducing formation of constitutive stress granules will stabilize mRNA as well.

## 1. Introduction

During its life span, the cell encounters various stresses and needs to regulate genome-wide gene expression to survive. An example of such regulation occurs in response to glucose starvation in the yeast *Saccharomyces cerevisiae*. Lack of glucose triggers general translational repression and downregulation of a number of transcripts [1-3]. At least some mRNAs abundant in glucose-replete conditions are targeted to P bodies [3,4]. At the same time, some transcripts are transcriptionally upregulated and are either directed to stress granules and P bodies (such as mRNAs involved in glucose metabolism) or diffusely localized in cytoplasm and actively translated (such as mRNAs involved in stress response) [3]. Such differential regulation allows the cell to adequately respond to the stress conditions and to sequester mRNAs that will be in high demand once the stress is relieved.

Stress granules and P bodies are induced by a number of stress factors and are composed of several proteins and mRNAs whose translation is attenuated during the stress (reviewed in [5,6]). Depending on the source of stress, formation of stress granules and P bodies may be driven by distinct mechanisms.

Under glucose starvation, formation of P bodies precedes assembly of stress granules; stress granules are primarily formed on pre-existing P bodies and their assembly depends on P body formation [2]. Phosphorylation of Dcp2 by Ste20 protein kinase and subsequent targeting of phosphorylated Dcp2 to P bodies is required for the assembly of stress granules, whereas P bodies are formed independently of Dcp2 phosphorylation and stress granules assembly [2,7]. Assembly of stress granules and P bodies relies on the function of several proteins, some of which contribute to the assembly of both these structures [2,8]. The most drastic phenotype of decreased percentage of cells with stress granules is observed in *pub1Δ* strain, whereas formation of P bodies in this strain is not affected [2].

Stress granules formed in response to sodium azide treatment differ from the granules formed in glucose-deprived cells in terms of protein composition as well as simultaneous formation with P bodies in generally non-overlapping pattern and independent manner [9]. Several protein factors that significantly contribute to the stress granules assembly in glucose deprived cells (including Pub1) have only minor or no effect upon sodium azide treatment. At the same time, requirements for protein factors in P body assembly are more conserved for these two stresses [9].

Stress granules induced by robust heat shock (46°C) are similar to stress granules formed upon sodium azide treatment and distinct from those induced by glucose starvation in terms of protein composition and Pub1-independent assembly [2,9,10]. P body components Dcp2 and Dhh1 co-localize with stress granules induced by robust heat shock; however, in certain circumstances, Dcp2 foci may dissociate from the stress granules [10].

Oxidative and osmotic stresses induce P bodies but no or very few stress granules [2,11]. P bodies formed under osmotic stress are more abundant relative to the ones formed under glucose starvation, and in this high abundance resemble P bodies formed in secretory pathway mutants [11]. However, more detailed analysis has shown that in these two cases the highly abundant P bodies are formed by different mechanisms in terms of requirement for calmodulin and P body components Pat1 and Scd6 [11].

High cell density induces both stress granules and P bodies. However, these structures are formed at different points of time and generally do not co-localize [12]. Under these conditions, the absence of Pub1 has a drastic effect on P body formation, whereas stress granules are formed at the wild-type level [12]. Examining a number of stress factors the authors also demonstrated that the formation of P bodies requires Pat1 and depends on phosphorylation of Pat1 by cAMP-dependent protein kinase (PKA), whereas formation of stress granules occurs independently of this mechanism [12,13].

Although different mechanisms are involved in the formation of stress granules and P bodies in distinct stress conditions, several stresses such as hyperosmolarity, robust heat shock, glucose deprivation, high cell density and sugar-induced osmotic stress lead to stabilization of multiple yeast mRNAs [14-16]. Genome-wide analysis have demonstrated global stabilization of yeast mRNAs during severe osmotic stress [17]. Stabilization of a number of transcripts is suggested under shift from glucose to galactose [18] and during oxidative stress [19]. The primarily mechanism of the stabilization is inhibition of deadenylation, which occurs either prior to or at the step of poly(A) shortening [14,15].

In this study, we address the possibility to accumulate mRNA with mammalian codon bias, which we termed the model RNA in yeast mutants with constitutive stress granules or elevated P bodies [20]. We rationalized that at least in some of the mutants the mRNA stabilization mechanism described above will be activated and the mRNA will be protected from degradation. In the light of the growing field of mRNA vaccines, the possibility of such an accumulation could represent a starting point for further work in production of specific capped and polyadenylated vaccine mRNAs as an alternative to the *in vitro* transcription, capping and following HPLC or FPLC purification required to remove double-stranded RNA contaminants (reviewed in [21]). Accumulated mRNA could also be used for *in vivo* assembly of heterologous RNA-protein complexes and for efficient recombinant protein expression in case an inducible way of its exit from silenced state to translation is established.

We demonstrate that the model RNA accumulates in *cho2Δ, rlf2Δ, rpl42aΔ* and *rtc2Δ* mutants, which are known to form constitutive stress granules overlapping with P bodies [20]. We further suggest that the accumulation occurs due to the protection of the model RNA from 5′ to 3′ degradation, as the combination of *dcp2-7, cho2Δ* and *rtc2Δ* mutations in one strain does not result in higher extent of the accumulation than in a single mutant. Our data suggest that the formation of constitutive stress granules and accumulation of the model RNA occur independently as the extent of the accumulation does not correlate with the extent of stress granules phenotype and in some mutants with constitutive stress granules the model RNA does not accumulate and may be downregulated. Moreover, there is no obvious accumulation of stress granule factors Pab1 and Pub1 at the sites of accumulation of the model RNA. We suggest that formation of stress granules in the mutants [20] is driven by distinct mechanisms, some of which do not simultaneously induce mRNA stabilization at least in the context of a certain mRNA.

## 2. Results

### 2.1. The model RNA is not downregulated due to its suboptimal codon bias

To investigate the possibility of accumulating mRNA with mammalian codon bias in yeast, we constructed vectors containing *EGFP* (mammalian codon-optimized enhanced green fluorescent protein) placed after mammalian consensus Kozak sequence. The *EGFP* open reading frame (ORF) sequence was followed by three MS2 binding sites and *CYC1* 3′UTR (see materials and methods for details). The introduction of the three MS2 stem-loops aimed to provide the possibility for a subsequent affinity purification of the accumulated mRNA or mRNA-protein complexes. We termed this transcript “the model RNA” and expressed it in yeast either under control of *GPD* promoter or in Tet-Off system.

Previous studies reported that mRNAs with non-optimal codon bias are destabilized in yeast even if the stretches of the rare codons are short [22,23]. Therefore, we first tested whether the model RNA had suboptimal codon sequences that could affect its steady state level. To this end, we compared codon optimality of the model RNA and *EGFP* mRNA optimized for translation in yeast (*yEGFP*) and known to be efficiently translated [24]. To each codon of *EGFP* and *yEGFP*, we assigned codon stabilization coefficient (CSC) determined in recent genome-wide study in yeast [22]. The CSC ranged from 0.3 (most optimal codons) to −0.3 (most non-optimal codons) and was color-coded from green to red, respectively. We found that the ORF of *EGFP* bore several stretches of amino acid codons suboptimal for expression in yeast (Figure 1A), and next, we tested whether this suboptimal codon bias would compromise the EGFP expression and abundance of the model RNA. To this end, the construct was generated where *EGFP* ORF in the model RNA was replaced by *yEGFP* ORF. Because for optimal translation *yEGFP* was placed under control of yeast consensus Kozak sequence, a control construct expressing *EGFP* under control of yeast consensus Kozak sequence was used to exclude the effect of Kozak sequence on the mRNA levels. As expected from the suboptimal codon bias, EGFP signal in cells expressing the model RNA was significantly lower than in cells expressing *yEGFP*, presumably due to less efficient translation. Yeast consensus Kozak sequence had only a minor effect on the EGFP expression (Figure 1B). At the same time, we found that steady state levels of the three transcripts were similar (Figure 1C). Thus, codon suboptimality of the model RNA did not significantly affect its abundance in yeast relative to codon-optimized transcript.

**Figure 1.**
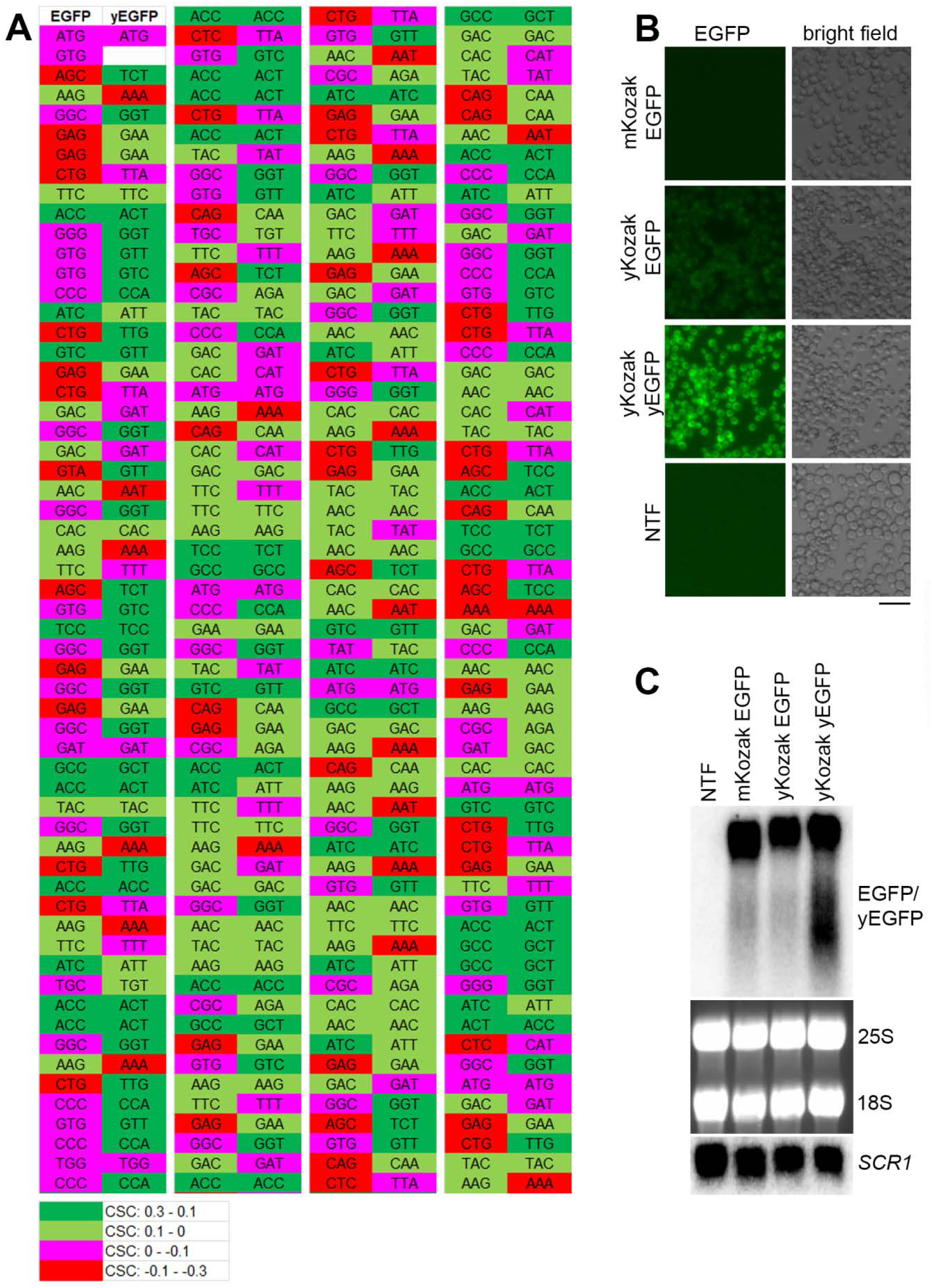
The steady state level of the model RNA is unaffected by its suboptimal codon bias. (A): comparison of codon optimality for expression in yeast between *EGFP* and *yEGFP*. CSC was attributed to each amino acid codon according to [22] and color-coding was added, as indicated in the lower panel. For image convenience the open reading frame (ORF) of *EGFP* and *yEGFP* was divided in four regions depicted from left to right and separated by vertical white line. (B): EGFP fluorescence (EGFP) was detected by live-cell imaging in strains expressing the model RNA (mKozak EGFP), the *EGFP* RNA flanked with yeast optimized Kozak sequence (yKozak EGFP) and the *yEGFP* RNA flanked with yeast optimized Kozak sequence (yKozak yEGFP). All the constructs above were integrated into *GPD* locus and expressed under control of *GPD* promoter. NTF indicates non-transformed cells (negative control for unspecific signal). Scale bar is 20 µm. (C): Northern blot analysis on total RNA isolated from the strains used for live-cell imaging in (B). The probe for Northern detection was located downstream of *EGFP/yEGFP* ORF in the region identical for all three RNAs. *SCR1*, 25S and 18S RNAs were used as endogenous controls. *SCR1* was detected by Northern blot and ribosomal RNA was detected by ethidium bromide staining.

### 2.2. The model RNA accumulates in yeast mutants with constitutive stress granules

A recent study identified a number of non-essential yeast gene deletion mutants with elevated size and number of P bodies and constitutive stress granules in the absence of stress [20]. As at least some of the mutants contained poly (A)^+^ mRNA in these cytoplasmic structures [25] and stabilization of a number of yeast mRNAs was reported under stress conditions (see introduction), we rationalized that at least some of these mutants may accumulate the model RNA, presumably by sequestering it in stress granules or P bodies. Therefore, we performed screening of 13 mutants with the most pronounced phenotype of constitutive stress granules and/or elevated P bodies [20]. In the screening, we also included the *atg15Δ* mutant, which showed pronounced accumulation of stress granule and P body markers in an intravacuolar compartment [20], and two *dcp2* mutants (*dcp2-7* and *dcp2Δ*) as *dcp2Δ* mutant forms numerous P bodies in the absence of stress [7] and *dcp2-7* mutation induces overlapping P bodies and stress granules at the restrictive temperature [20]. We expressed the model RNA on episomal plasmid in Tet-Off system, and we tested its steady state levels in the mutants by Northern blot.

Among the mutants that were shown to form constitutive stress granules overlapping with P-bodies [20], we found elevated levels of the model RNA in *dcp2-7* and *xrn1Δ* mutants (Figure 2 and Supplementary File 1: Table S1), which is expected as Dcp2 and Xrn1 act in mRNA decapping and 5′ to 3′ degradation generally considered as a major pathway of mRNA surveillance and decay in yeast (reviewed in [26]). In addition, accumulation of the model RNA was systematically observed in gene deletion mutants of four novel factors: *RPL42A, CHO2, RLF2* and *RTC2* (Figure 2 and Supplementary File 1: Table S1). All the four gene deletion mutants were shown to form constitutive stress granules overlapping with P bodies and two of them (*rpl42aΔ* and *rlf2Δ*) also formed enlarged P bodies [20]. Interestingly, the extent of accumulation of the model RNA in these mutants did not correlate with the extent of constitutive stress granules phenotype. For example, constitutive stress granules were found in approximately 30% in *rtc2Δ* and *cho2Δ* mutant cells and in 8% of *rpl42aΔ* mutant cells, whereas the accumulation of the model RNA in these mutants was similar (Supplementary File 1: Table S1). Consistent with this observation, seven other mutants with pronounced constitutive stress granules, six of which formed stress granules overlapping with P bodies and five had elevated P bodies, did not show systematic accumulation of the model RNA: the steady state level of the model RNA varied between the experiments and, in some cases, was downregulated (Supplementary File 1: Table S1). The mutant with stress granules distinct from P bodies (*mft1Δ*) systematically showed downregulation of the model RNA and in *atg15Δ* mutant the steady state level of the model RNA varied between the experiments, with a tendency to downregulation (Figure 2 and Supplementary File 1: Table S1).

**Figure 2.**
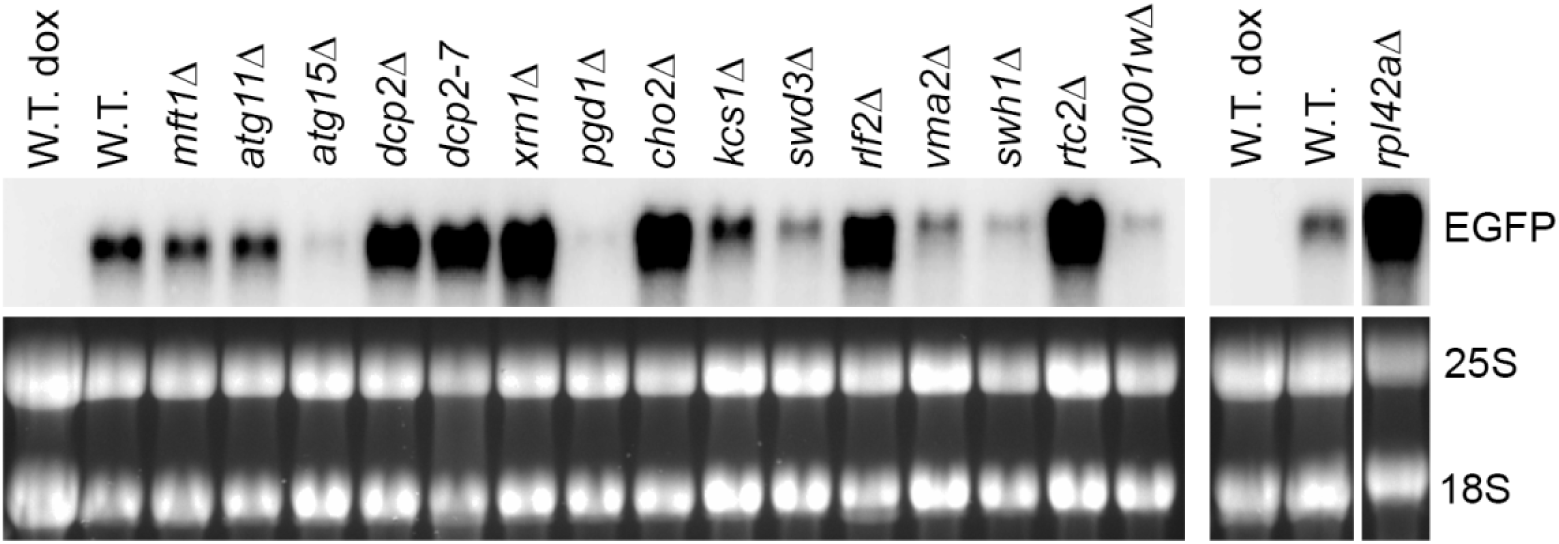
The model RNA accumulates in mutants with constitutive stress granules. Northern blot analysis was performed on total RNA from wild-type strain (W.T., BY4741) and mutants (indicated above each line) transformed with episomal plasmid expressing the model RNA in Tet-Off system. The model RNA was detected with the probe to *EGFP*. Ribosomal RNA (25S and 18S) was visualized by ethidium bromide staining and served as endogenous control. RNA from uninduced cells (dox) served as control for unspecific hybridization. Wide white vertical line indicates that the samples were run on two different gels. Narrow white vertical line indicates that the samples were run on the same gel, but some lines between left and right part were removed. Full-length gel and blot for the cropped images are shown in Supplementary File 1: Figure S3A.

In summary, the data demonstrate that formation of the cytoplasmic bodies in the mutants above does not necessarily lead to the accumulation of the model RNA and suggests that first, distinct mechanisms induce and regulate assembly of the cytoplasmic bodies in the mutants, and, second, that accumulation of the model RNA may occur independently from stress granules formation.

### 2.3. Accumulation of the model RNA in rtc2Δ and cho2Δ mutants occurs due to compromised path linked to the activity of Dcp2

Next, we investigated the possible mechanisms of accumulation of the model RNA. To this end, we analyzed expression of the model RNA in Tet-Off system in wild-type and mutant cells by fluorescent *in situ* hybridization (FISH). We did not observe any FISH signal in the presence of doxycycline, which, first, argues that the signal is specific to the location of the model RNA, and, second, shows that the leakage from the promoter in the absence of induction is extremely low or absent (Supplementary File 1: Figure S1A). Upon induction in the wild-type cells (BY4741), the model RNA is expressed at different levels within the cell population. The uneven expression occurs due to bursts in mRNA transcription from *TetO7* promoter [27,28]. Such transcriptional bursts were shown to regulate mRNA expression in a subset of genes; however, this phenomenon is not a general path for the regulation of gene expression in yeast [29,30].

A group of cells with very high model RNA expression level could be unequivocally distinguished as the FISH signal in these cells is several folds higher than in the rest of the population. When imaged with the same acquisition settings, the highly expressing cells show saturated FISH signal, whereas medium to low expressing cells are reasonably bright. Once the acquisition settings are adjusted to highly expressing cells, the cells with medium to low expression are no longer visible (Supplementary File 1: Figures S1A and B). This drastic difference in the expression levels is neither a specific feature of a particular strain nor an artefact of an acquisition method: Supplementary File 1: Figures S1A and B show reproducibility of this observation for BY4741 and *PAB1-GFP* strains where FISH signal was acquired with epifluorescence and confocal microscopy respectively.

We first asked if the accumulation of the model RNA in the mutants is linked to changes in the above phenomenon, which would lead to an evenly high expression. Under the screening conditions (episomal plasmid in Tet-Off system) the model RNA is expressed in approximately 50% of the wild-type cells with 16% of the total number of cells expressing it at the very high level (Figure 3A and Supplementary File 1: Table S2). In the mutants (*cho2Δ, rpl42aΔ* and *rtc2Δ*) that accumulate the model RNA, the expression remained uneven within the population, but the number of expressing cells in the population increased to 70–78%. For *cho2Δ* and *rpl42aΔ* mutants, the number of highly expressing cells was also higher than in the wild type: 27% and 40% respectively versus 16% in the wild type. This uneven increase of expression within the cell population was similar to the phenotype of *dcp2-7* mutant in which 25% of the cells expressed the model RNA at the high level (Figure 3A and Supplementary File 1: Table S2).

**Figure 3.**
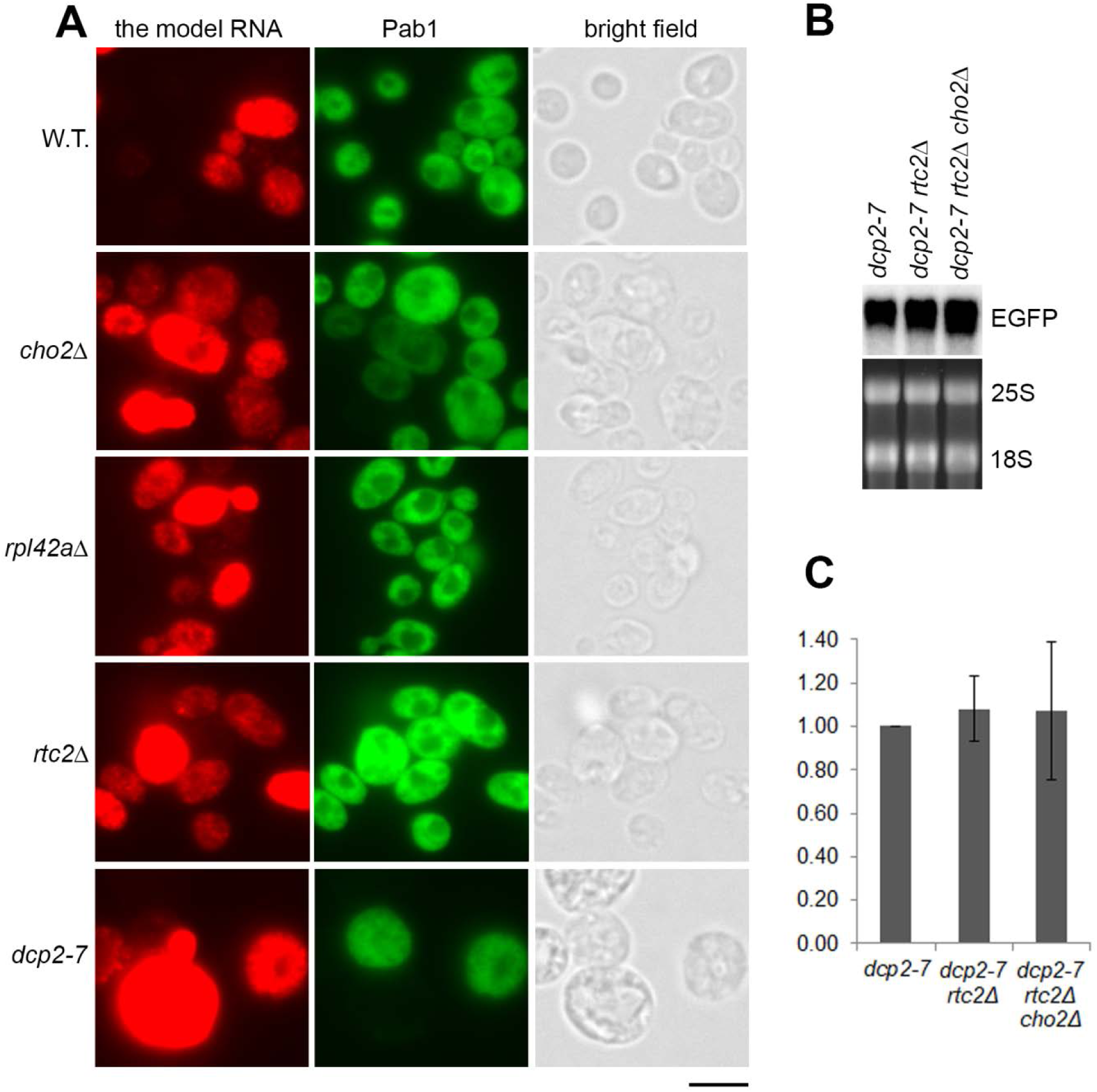
Accumulation of the model RNA in *rtc2Δ* and *cho2Δ* mutants is linked to compromised 5′ to 3′ degradation path. (A): The model RNA was expressed in wild-type (BY4741) and mutant strains (indicated on the left), as in Figure 2. Expression of the model RNA and Pab1 was then simultaneously detected by FISH and immunofluorescence, respectively (FISH-IF). Cells were examined by epifluorescence microscopy and manually counted. Scale bar is 5 µm. (B and C): Combining of *dcp2-7, rtc2Δ* and *cho2Δ* mutations in one strain does not lead to further accumulation of the model RNA. The model RNA was expressed as above in single, double and triple mutants (indicated above each line) and its steady state levels were detected by Northern blot (B), quantified using phosphoimager and expressed as fold increase relative to the levels in *dcp2-7* mutant (C). Ribosomal RNA (25S and 18S) was visualized by ethidium bromide staining and served as endogenous control. Quantitation in (C) represents data from three experiments.

The similarity in accumulation phenotype between the constitutive stress granules mutants and *dcp2-7* strain prompted us to ask whether the accumulation of the model RNA in the mutants is linked to the same path. To test this hypothesis, we deleted *RTC2* gene in *dcp2-7* background and subsequently deleted *CHO2* gene in the double mutant. Northern blot analyses have shown that combining *dcp2-7* and *rtc2Δ* mutations as well as *dcp2-7, rtc2Δ* and *cho2Δ* mutations in one strain did not lead to a further increase in the steady state levels of the model RNA (Figure. 3B, C). Two independent clones of each *dcp2-7 rtc2Δ* and *dcp2-7 rtc2Δ cho2Δ* mutants were tested and furnished the same results (Supplementary File 1: Figure S1C). Thus, we concluded that accumulation of the model RNA in *rtc2Δ* and *cho2Δ* mutants occurred in the same path as in *dcp2-7* mutant.

We also found that in the wild-type strain, 96% cells expressed Pab1, whereas only 24% cells in *dcp2-7* mutant expressed Pab1 under restrictive temperature (Figure 3A and Supplementary File 1: Table S2). This observation is in line with a report demonstrating downregulation of Pab1 mRNA in *dcp2Δ* strain to approximately 25% of the wild-type level [31]. As Pab1 is known to be an inhibitor of mRNA decapping [32], this downregulation may reflect a regulatory mechanism that occurs when Dcp2 is missing or inactive. At the same time, we found that this mechanism is not activated in the mutants that accumulate the model RNA: Pab1 was expressed in these mutants at the wild-type level (Figure. 3A and Supplementary File 1: Table S2).

### 2.4. The model RNA forms cytoplasmic granules distinct from stress granules and P bodies

To further investigate the possible mechanisms of accumulation of the model RNA, we asked whether it can be targeted to stress granules or P bodies and thus be protected from 5′ to 3′ surveillance.

Upon induction in the Tet-Off system in the wild-type cells (BY4741), the model RNA forms granular pattern in the cytoplasm (Figure 4A). As it was mentioned above, approximately 50% of the wild-type cells expressed the model RNA.

**Figure 4.**
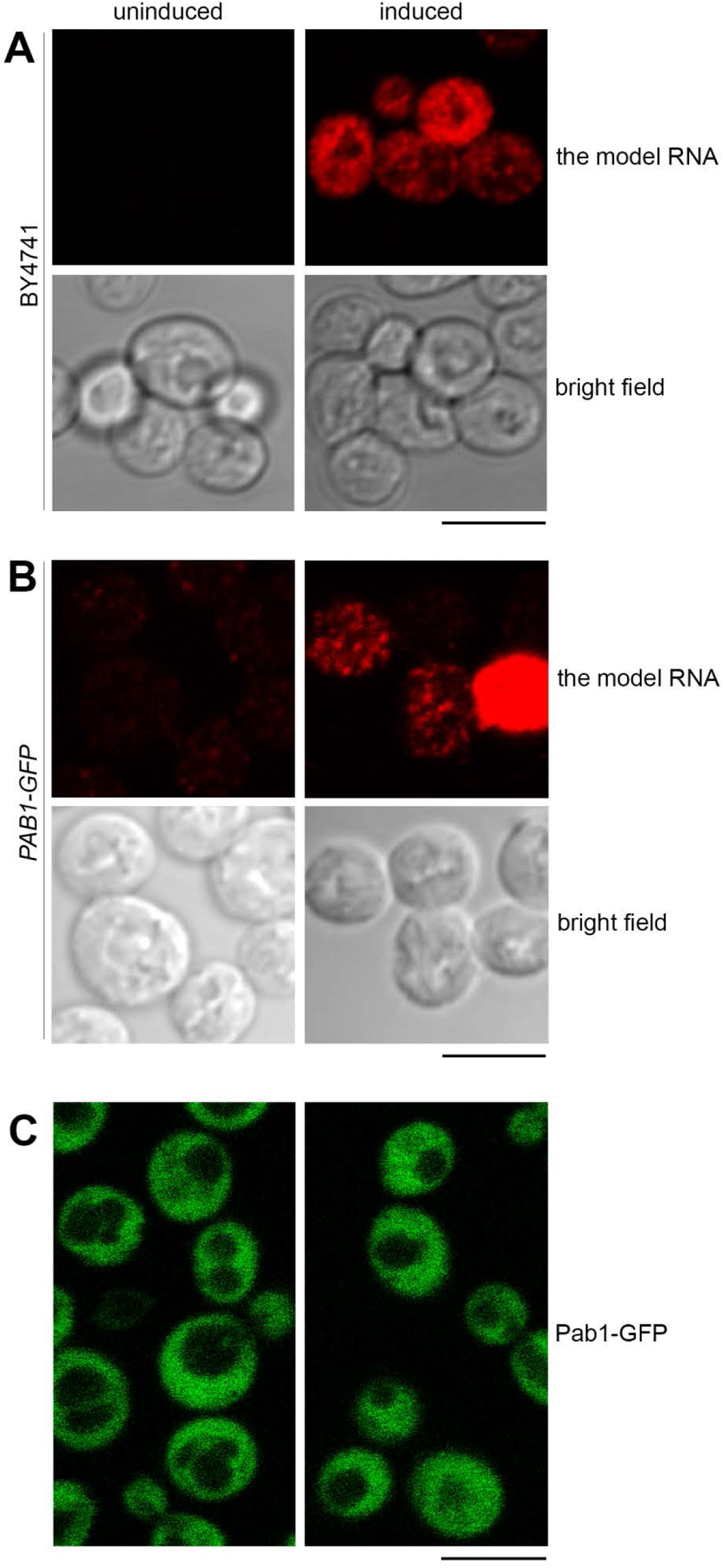
The model RNA is located to cytoplasmic granules distinct from stress granules. (A and B): Expression of the model RNA was visualized by FISH in wild-type (BY4741), (A) and in *PAB1-GFP* (B) strains and showed similar pattern of cytoplasmic granules formed in approximately 50% of cells. (C): Live-cell imaging of *PAB1-GFP* strain expressing the model RNA did not detect formation of stress granules. In all panels the model RNA was expressed as in Figure 2, and uninduced cells were used to control for unspecific FISH signal (A and B) and for appearance of stress granules unrelated to the expression of the model RNA (C). Scale bar is 5 µm.

We first asked whether this granular pattern could represent targeting of the model RNA to the stress granules. We used Pab1 as a conventional marker for stress granules and expressed the model RNA as above in *PAB1-GFP* strain where chromosomal copy of the *PAB1* gene is tagged with *GFP*. In a manner similar to BY4741, approximately 50% of *PAB1-GFP* cells expressed the model RNA with granular cytoplasmic pattern and minimal signal in the absence of induction (Figure 4B, Supplementary File 1: Figure. S1B and data not shown). At the same time, we did not observe targeting of Pab1-GFP to cytoplasmic bodies within this cell population examined by live-cell imaging (Figure 4C). Thus, we consider it unlikely that expression of the model RNA could generate formation of stress granules.

We next asked if the granular cytoplasmic pattern formed by the model RNA could be linked to its extensive targeting to P bodies. To test this hypothesis, we used *EDC3-GFP* strain where endogenous *EDC3* gene is tagged with *GFP*. Under normal conditions, the majority of untransformed *EDC3-GFP* cells form one to three small P bodies per cell (Figure 5A). Upon induction, approximately 60% of *EDC3-GFP* cells express the model RNA with minimal background signal (Figure 5C and data not shown). However, the induction does not lead to the increase in size and/or number of P bodies within the cell population (Figure 5B and Supplementary File 1: Table S3). Using simultaneous labeling by FISH and immunofluorescence (FISH-IF), we demonstrated that while multiple granules of the model RNA are distributed throughout the cytoplasm of *EDC3-GFP* cells, P bodies maintain low abundance and generally do not co-localize with the model RNA, although they may locate in close vicinity of one or two granules (Figure 5C).

**Figure 5.**
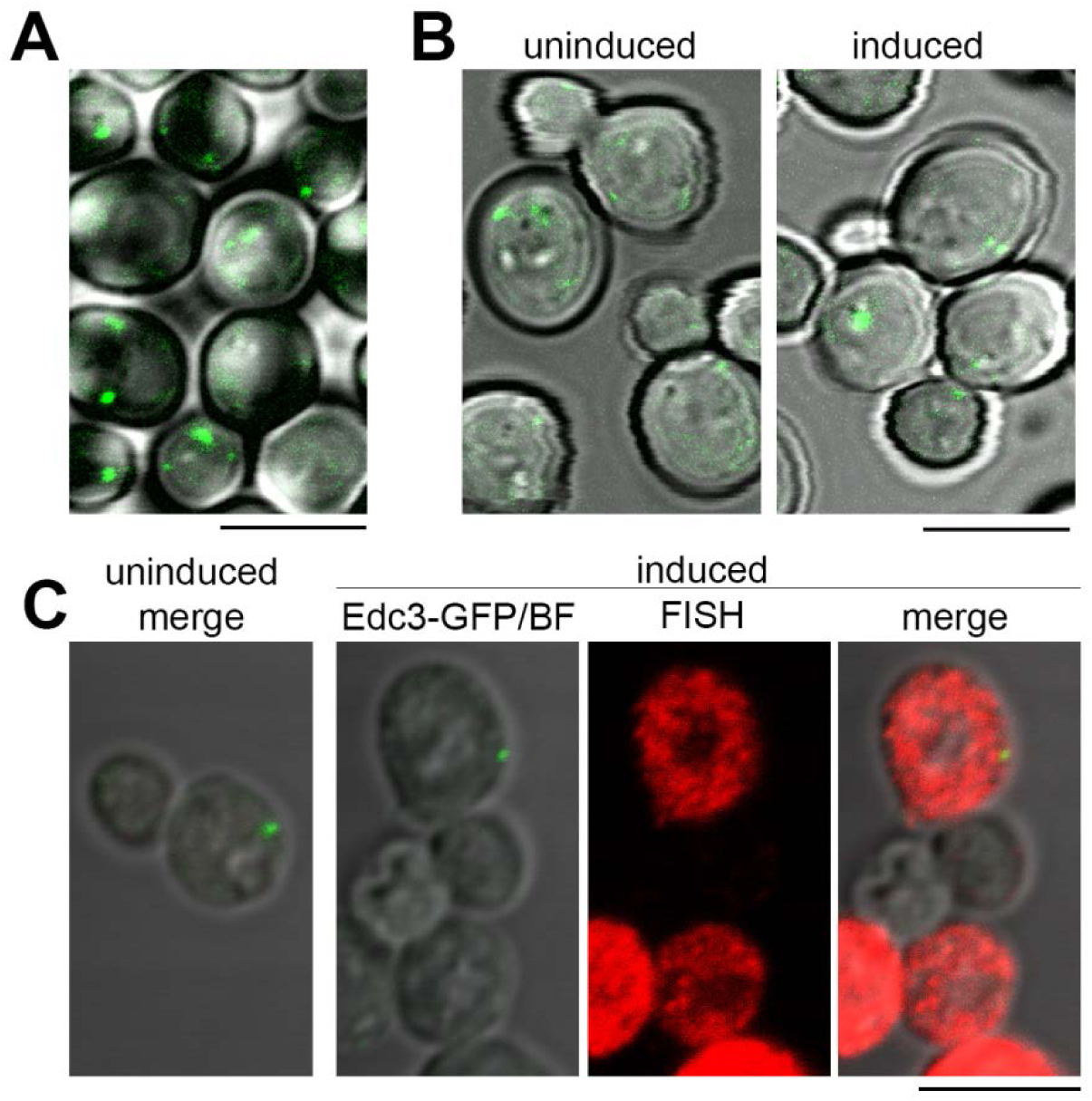
Expression of the model RNA does not induce formation of P bodies. (A): Live-cell imaging of non-transformed *EDC3-GFP* strain. Generally, under normal conditions, between 1 and 3 small P bodies per cell are detected. (B and C): The model RNA was expressed in *EDC3-GFP* strain as in Figure 2. Uninduced cells were used to control for P bodies unrelated to the expression of the model RNA and for unspecific FISH signal. (B): Live-cell imaging of *EDC3-GFP* strain. Expression of the model RNA leads to the increase in neither the size nor the number of P bodies per cell. (C): FISH-IF of the model RNA and Edc3-GFP showed that the granules of the model RNA are distinct from P bodies, although some of them may be located in the vicinity of each other. In (A) and (B), overlay of Edc3-GFP signal and bright field is shown. In (C), “FISH” corresponds to FISH detection of the model RNA, “Edc3-GFP/BF” corresponds to overlay of bright field and Edc3-GFP signal and “merge” corresponds to overlay of bright field, FISH and Edc3-GFP labeling. Scale bar is 5 µm.

Thus, we concluded that in wild-type cells, the model RNA forms cytoplasmic granules distinct from stress granules and P bodies.

We next asked if accumulation of the model RNA in the mutants could be linked to its targeting to stress granules. To this end, we analyzed distribution of the model RNA in the mutants using FISH-IF with two antibodies that recognize stress granules markers Pab1 and Pub1 [33]. As with the wild type, we did not observe any targeting of Pab1 to cytoplasmic granules in the mutants that express the model RNA (Figure 6, left panel). Surprisingly, we did not find stress granules in mutant cells that do not express the model RNA (data not shown). The total number of examined cells is indicated in Supplementary File 1: Table S2.

**Figure 6.**
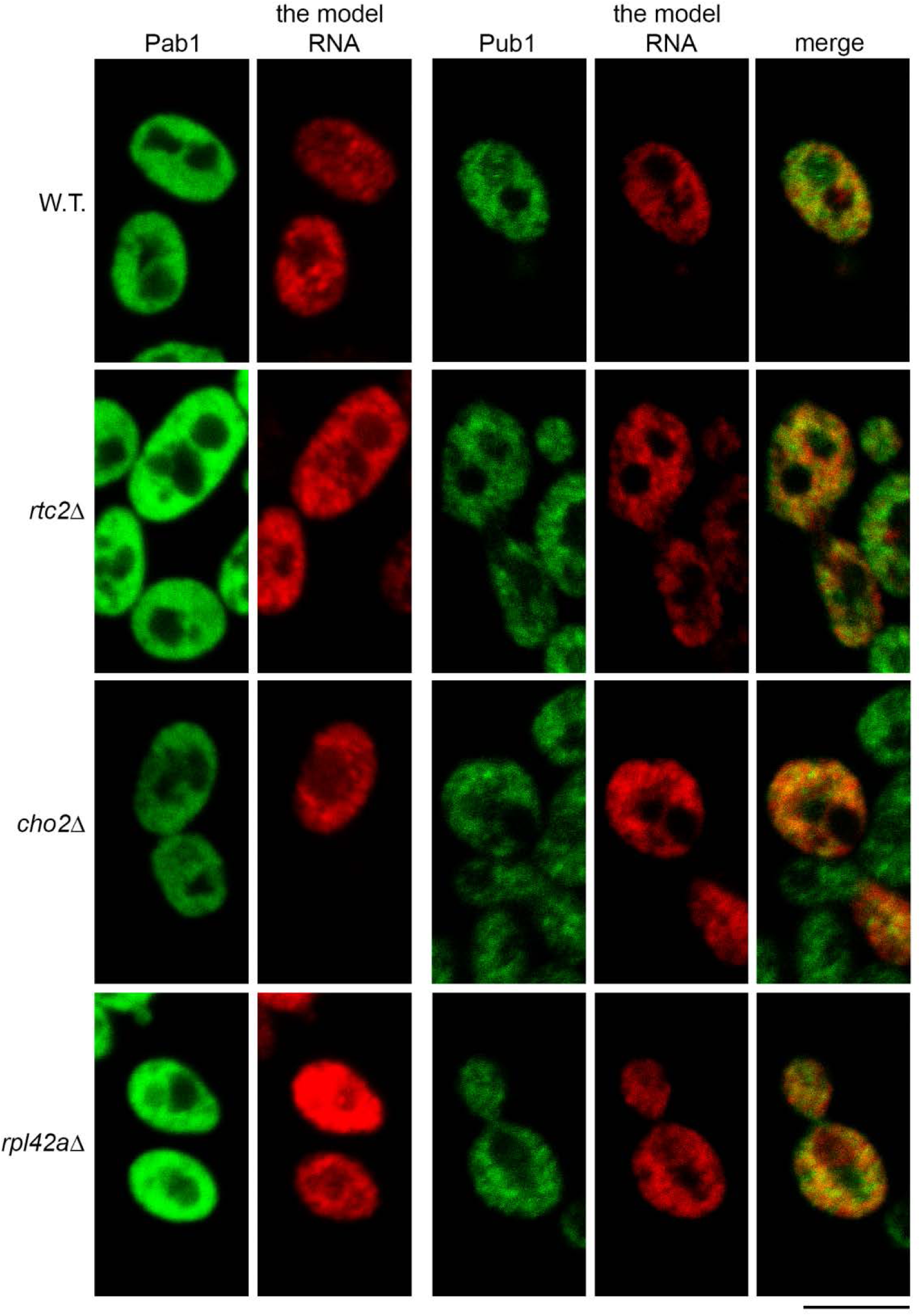
Accumulation of the model RNA in the mutants is not linked to its targeting to stress granules. FISH-IF of the model RNA (expressed as in Figure 2) and stress granule markers Pab1 (left panel) and Pub1 (right panel) in wild-type (BY4741, W.T) and mutant strains as indicated on the left. Scale bar is 5 µm.

As with the previous report [33], we found that Pub1 formed granular pattern in immunofluorescent labeling of wild-type cells. However, these granules did not generally co-localize with the granules of the model RNA, although some partial co-localization may occur (Figure 6, right panel). In *cho2Δ, rpl42aΔ* and *rtc2Δ* mutants, which accumulate the model RNA, the granular pattern of Pub1 was similar to that of the wild type, without obvious formation of distinct stress granules. As in the wild type, partial co-localization of Pub1 granules and the model RNA was observed. However, there was no more obvious extensive co-localization than in the wild type cells (Figure 6, right panel). Thus, our data do not support the idea that accumulation of the model RNA is linked to its targeting to the stress granules in the mutants.

### 2.5 Accumulation of the model RNA and defects in endoplasmic reticulum

A previous study demonstrated sequestration of *WSC2* mRNA tagged with MS2 loops in aberrant structures of endoplasmic reticulum (ER) in *cho2Δ* mutant [34]. Therefore, we asked whether ER in the mutants that accumulate the model RNA could have abnormalities that would contribute to the accumulation. To detect ER, we expressed RFP-ER, a reporter in which transmembrane domain of ER protein Scs2 is fused to the tandem repeat of dimeric DsRed [35]. In a manner similar to the previous report [34], we found that in *cho2Δ* mutant RFP-ER was detected in aberrant aggregate-like structures and cortical ER was less intensively stained than in the wild type (Figure 7). In *rtc2Δ* mutant, the aggregate-like structures were also found, and the staining of cortical ER was distinct from the wild type (Figure 7). At the same time, *rpl42aΔ* mutant showed normal outlines of ER, although systematically less intensively stained than in the wild type (Figure 7). Induction of the model RNA led to down-regulation of RFP-ER almost to the background both in wild-type strain and in the mutants (Supplementary File 1: Figure S2). Therefore, we suggest a possible link between expression of the model RNA, its accumulation at least in some mutants and ER morphology. However, the mechanism is unlikely to be identical to the accumulation of *WSC2* mRNA and needs to be studied further.

**Figure 7.**
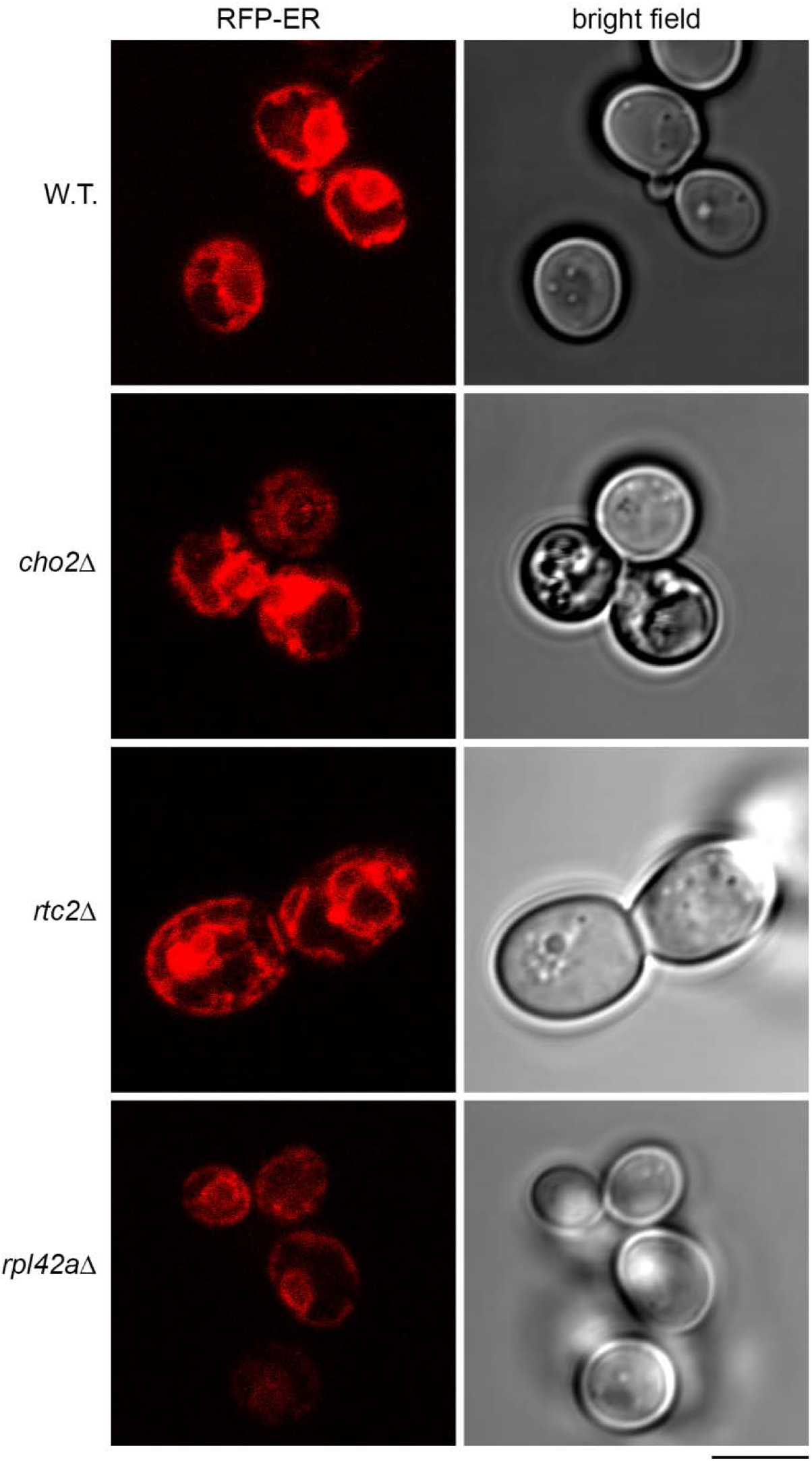
Endoplasmic reticulum of the mutants accumulating the model RNA shows deviations from the wild type. Live-cell imaging of wild-type (BY4741, W.T.) and mutant strains (indicated on the left) transformed with the plasmid expressing RFP-ER. Scale bar is 5 µm.

## 3. Discussion

In this study, we report the possibility of accumulating an mRNA with mammalian codon bias in yeast mutants known to form constitutive stress granules in the absence of stress with up to an 8-fold increase relative to the wild type (Figure 2 and Supplementary File 1: Table S1). Stabilization of mRNAs under stress conditions was reported in several studies and relies on protection from deadenylation (see introduction). Deadenylation is the first step in cytoplasmic mRNA degradation in *S. cerevisiae*. Following deadenylation, mRNA is either degraded from 3′ to 5′ by the exosome or decapped by the Dcp1/Dcp2 decapping enzyme and degraded from 5′ to 3′ by the exonuclease Xrn1 (reviewed in [26]). Our data show that, at least in two mutants, accumulation of the model RNA occurs in the same path as in *dcp2-7* mutant, suggesting that the mechanism may be linked to compromised 5′ to 3′ surveillance (Figure 3). At the same time, the growth of the mutants accumulating the model RNA is not severely affected as it is known for *dcp2* mutant strains (data not shown), and the expression of Pab1 in these mutants is not compromised in contrast to its downregulation in *dcp2* mutants (see results). These observations bring a certain doubt that Dcp2 function is generally affected in the mutants which accumulate the model RNA. In summary, all the data above fit well with the view that the mechanism of mRNA stabilization under stress conditions is activated in the mutants with constitutive stress granules, which accumulate the model RNA, and that this mechanism relies on protection of mRNA from the main degradation path.

A possible mechanism of accumulation would be sequestration of the model RNA, so that it is no longer accessible to the decapping complex. Our data, however, do not support the view that the model RNA is sequestered in stress granules. First, expression of the model RNA in wild-type cells does not lead to the formation of stress granules or P bodies as judged by localization of endogenous Pab1 and Edc3 tagged with GFP (Figures 4, 5); thus, the data are incompatible with the idea that the model RNA is normally targeted to stress granules and accumulates in mutants due to the promotion of stress granules formation. Second, there is no absolute correlation between the increase in total amount of the model RNA in mutants that accumulate it and the extent of the reported elevated stress granule phenotype in these mutants (Supplementary File 1: Table S1). Finally, our FISH-IF experiments have demonstrated that two stress granules markers, Pab1 and Pub1, do not accumulate at the sites where the model RNA is detected in the mutants that accumulate it (Figure 6). Thus, our data are incompatible with the idea that only some amount of the model RNA is sequestered in stress granules, thus leading to the accumulation.

Surprisingly, in our hands, FISH-IF experiments with both Pab1 and Pub1 failed to identify stress granules among hundreds of cells in the mutants that accumulate the model RNA. Possible reasons for the discrepancy with the reported data [20] may be on distinct detection systems for the stress granules. First, the stress granules detected by live cell imaging in these mutants may be generally quite unstable and easily lost during the FISH-IF procedure. Second, Pab1 and Pub1 may not be efficiently targeted to the stress granules in the mutants and/or be associated with them in a very dynamic manner. At the same time, behavior of GFP-tagged Pab1 and Pub1 introduced on the plasmid may be slightly distinct from endogenous proteins in terms of more efficient targeting and longer residence time in stress granules in these mutants. The fact that only a portion of cells forms stress granules in the mutants [20] supports both the possibilities above.

Another possible mechanism for accumulation of the model RNA could be compromised activity of the decapping complex on a subset of mRNAs. Post-translation modifications of Dcp2 and/or other components of 5′ to 3′ surveillance machinery could provide a possible link between the assembly of cytoplasmic bodies and selective stabilization of mRNAs. Under glucose starvation, phosphorylation of yeast Dcp2 is required for the assembly of stress granules, stabilizing a subset of mRNAs at the same time [7]. Phosphorylation of human Dcp1a (hDcp1a) regulates assembly of P bodies and stabilizes certain mRNAs [36]. During mitosis, when the bulk of the cellular RNAs need to be protected from degradation, hDcp1a is also phosphorylated, these phosphorylation events playing a role in its association with P bodies [37]. We hypothesize that constitutive stress granules in the mutants may be formed in a path where some components of the decapping machinery are post-translationally modified so that the complex is less active on certain substrates and the loss in activity could be sufficient to accumulate the model RNA. At the same time, such post-translational modifications may trigger assembly of constitutive stress granules.

Several distinct mechanisms lead to the formation of stress granules under various stresses and the protein composition as well as the key assembly factors vary significantly depending on the source of stress and some other factors (see introduction). The term “stress granules” was also attributed to the cytoplasmic structures formed in the number of mutants in a genetic screen based on simultaneous targeting of Pab1 and Pub1 fused to GFP to these structures [20]. However, a more detailed investigation into stress granules formed in mutants of THO/TREX2 complex has demonstrated that the composition of these structures comprises of not only typical stress granules factors but also proteins so far not found in stress granules; moreover, assembly of stress granules induced by sodium azide was impaired in some of these mutants [25]. Our data demonstrates that formation of stress granules in the mutants screened in this study can be driven by distinct mechanisms as formation of stress granules *per se* does not lead to the accumulation of the model RNA. In several mutants with constitutive stress granules, the steady state levels of the model RNA varied between experiments, and in the case of *mft1Δ* mutant, where up to 33% cells form stress granules, the model RNA was systematically downregulated. Further investigations are required to identify the link between stress granules formed under stress conditions and in the mutants accumulating the model RNA as well as to reveal the mechanisms that coordinate stress granules assembly and RNA stabilization during stress. Our findings, however, suggest that these two paths could be uncoupled in certain mutants, which provides a platform for further studies.

## 4. Materials and Methods

### 4.1. Plasmids

RFP-ER in YCp50 was described [35]. We confirmed by sequencing that the RFP variant in this plasmid is DsRed.

All the oligonucleotides used in this study were purchased from Eurogentec, Belgium.

Three MS2 binding sites were generated in pGEM vector as described [38] in Prof. Maria Carmo-Fonseca’s lab (N. V. Kozlova, unpublished data). To generate the model RNA, the MS2 binding sites were amplified with primer pair 1 (Supplementary File 1: Table S4) and cloned into pEGFP-C1 (Clontech) as EcoRI/BamHI fragment. This cloning also introduced STOP codon in the end of *EGFP* open reading frame. The sequence containing *EGFP* and 3MS2 binding sites was subcloned as AgeI (blunted)/BamHI fragment into pGEM4Z/GFP/A64 [39] digested HincII/BamHI. The resulting plasmid was named pGEM4Z-EGFP-3MS2-A64. For expression of the model RNA in yeast, *GPD* promoter and *CYC1* terminator were amplified from genomic DNA with primer pairs 2 and 3 respectively (Supplementary File 1: Table S4). First, *CYC1* terminator was cloned into pFL26 [40] as BamHI/SacI fragment, yielding pFL26-CYC1. *GPD* promoter was cloned into pFL26-CYC1 as HindIII/SbfI fragment, yielding pFL26-GPD-CYC1. The sequence containing *EGFP* and MS2 binding sites was excised from pGEM4Z-EGFP-3MS2-A64 and inserted into pFL26-GPD-CYC1 as PstI/BamHI fragment. The resulting plasmid was named pYC4 and allowed to express the model RNA under control of *GPD* promoter. To express the model RNA in Tet-Off system, the EcoRI (blunted)/PstI fragment of pCM190 [41] bearing *TetO7* promoter and *tTA* was cloned into pYC4 replacing *GPD* promoter and yielding pYC6. The 2-microm replication origin of pCM190 was amplified with primer pair 4 (Supplementary File 1: Table S4) and inserted into pYC6 linearized by BspQI by gap repair [42]. The resulting plasmid was named pGR6 and used for episomal expression of the model RNA in Tet-Off system. High-fidelity enzymes were used for all PCR amplifications; the plasmids were sequenced and found to be identical to the expected.

### 4.2. Yeast strains and growth conditions

*PAB1-GFP* (12B2) *MATa his3Δ1 leu2Δ0 met15Δ0 ura3Δ0 PAB1-GFP::HIS3* and *EDC3-GFP* (13E1) *MATa his3Δ1 leu2Δ0 met15Δ0 ura3Δ0 EDC3-GFP::HIS3* were from Yeast-GFP clone collection from UCSF (Invitrogen). The identity of the GFP-tagging was confirmed by PCR. The *DCP2* mutants: *dcp2-7* (YAV747) *MATa his3Δ1 leu2Δ0 lys2Δ0 ura3Δ0 met15Δ0 dcp2-7::URA3* and *dcp2Δ* (YAV756) *MATa his3Δ1 leu2Δ0 lys2Δ0 ura3Δ0 dcp2Δ::NEO* were from Prof. Ambro Van Hoof. Non-essential gene deletion mutants in BY4741 background (*MATa his3Δ1 leu2Δ0 met15Δ0 ura3Δ0*) were from EUROSCARF. The identity of the gene deletions used in this study was confirmed by PCR with specific primers.

Two clones of *rtc2Δ* mutants were generated in *dcp2-7* background as described [43,44] using primer pair 5 (Supplementary File 1: Table S4) and the plasmid pFA6-kanMX4 [45]. The resulting genotype of the strains is as follows: *MATa his3Δ1 leu2Δ0 lys2Δ0 ura3Δ0 met15Δ0 dcp2-7::URA3 rtc2Δ::KanMX4*. One of these clones was used to generate *cho2Δ* mutants using primer pair 6 (Supplementary File 1: Table S4) and the plasmid pFA6a-hphNTI [44] as above. The resulting genotype of the strains is as follows: *MATa his3Δ1 leu2Δ0 lys2Δ0 ura3Δ0 met15Δ0 dcp2-7::URA3 rtc2Δ::KanMX4 cho2Δ::HphNTI*. Gene deletions were confirmed by PCR with specific primers.

To generate yeast strains expressing *yEGFP* and *EGFP* under control of yeast and mammalian consensus Kozak sequence, *yEGFP* was amplified from pKT128 [46] (Addgene) with primer pair 7, and *EGFP* was amplified from pYC4 with primer pairs 8 and 9, which introduced mammalian and yeast consensus Kozak sequence respectively (Supplementary File 1: Table S4). Approximately 100 ng of the PCR products were mixed with 250 ng of pYC4 digested SbfI/BspEI and transformed into W303 strain (*ade2-1 can1-100 his3-11,15 leu2-3,112 trp1-1 ura3-1*). The plasmids generated by gap repair were integrated into yeast genome. We confirmed by PCR that the three clones used in the Figure 1 were integrated into *GPD* locus. The integrated plasmids were recovered from genomic DNA as described [47] and the identity of the *EGFP* variant was confirmed by sequencing.

Yeast strains were grown according to standard procedures in complete synthetic medium composed according to nutrient requirements and plasmid selection. Doxycycline (dox) was added to the medium to the final concentration of 5 µg/ml to inhibit expression of the model RNA in Tet-Off system. In all experiments, cells were grown at 30°C, except for *dcp2-7* mutants, which were shifted to 37°C immediately after induction to assay for the phenotype. Frozen competent cells (FCC) protocol [48] was used for transformation of yeast strains with plasmids and PCR fragments.

### 4.3. Northern blot

Cells were grown overnight at low density on orbital shaker. For induction of the model RNA in Tet-Off system cells were washed four times with medium without dox and the optical density at 600 nm was measured. Ten ml of fresh medium was inoculated with 0.25 × 10^7^ cells/ml, cells were further grown for 6 hours and harvested.

For the Northern blot on Figure 1, cells were grown overnight as above, inoculated at 0.35×10^7^ cells/ml and grown to approximately 0.7×10^7^ cells/ml. Eight milliliters of each culture was harvested for Northern blot.

Total RNA was extracted as described [49] and used at 10 µg per line. Northern blot was performed according to standard procedures using lab modification of the method described by Rosen and co-authors [50]. Probes GTACAGCTCGTCCATGCCGAG and GGTCACCTTTGCTGACGCTGG were used to detect *EGFP* and *SCR1* RNAs respectively. Probe GATCCAGAGGCGGTACCG was used for simultaneous detection of *yEGFP* and *EGFP* mRNAs on Figure 1. The probe hybridizes to the RNA region located after MS2 binding sites which is common for all three constructs. Probes were end-labeled by kinase reaction and the intensity of the signal was quantified by phosphoimager.

### 4.4. FISH and immunofluorescence

Cells were grown overnight at low density on orbital shaker, washed as above, inoculated into fresh medium at 0.35 × 10^7^ cells/ml and harvested after 6 hours of induction. FISH was essentially performed as described [51] with custom synthesis probes (Eurogentec) GT*AGCCTTCGGGCAT*GGCGGACTTGAAGAAGT*CGTGCTGCTTCAT*GTGG, GGCT*GTTGTAGTTGT*ACTCCAGCTT*GTGCCCCAGGAT*GTTGCCGTCCT*CC, CT*GCACGCTGCCGT*CCTCGATGTT*GTGGCGGATCT*TGAAGTTCACCT*TGATG and AACT*CCAGCAGGACCAT*GTGATCGCGCTT*CTCGTTGGGGTCT*TTGCTCAG, where T* indicated amino-allyl thymidine. Probes were labeled with Cy3 or Cy5 Mono-Reactive Dye Pack (GE Healthcare).

For simultaneous detection by FISH and immunofluorescence (FISH-IF), FISH was performed as above, but instead of PBS the final wash was done in PBS supplemented with 0.05% Tween 20 (PBS-T). The samples were next incubated overnight in block solution (PBS-T, 3% BSA, 4 mM VRC), aspirated and incubated with primary antibodies in block solution for 1 hour at 37°C. After three washes for 5 minutes each with PBS-T, the samples were incubated with secondary antibodies in block solution for 45 minutes at room temperature, washed three times for 5 minutes each in PBS and mounted. Starting from the secondary antibody step, the samples were kept protected from light. Standard RNase-free practices were used throughout the protocol.

The signal from RFP-ER was enhanced with mouse monoclonal anti-DsRed antibody E8 (Santa Cruz Biotechnology) diluted 1/100. GFP was detected with Anti-GFP antibody (Roche) diluted 1/100. Endogenous Pab1 was detected with 1G1 antibody diluted 1/5000 and endogenous Pub1 was detected with 4C3 antibody diluted 1/100. Both the 1G1 and 4C3 antibodies [33] were from Prof. Maurice S. Swanson. Goat anti-mouse Alexa Fluor 488 and Alexa Fluor 568 conjugated antibodies (Invitrogen) were used in 1/200 dilution.

Images for the Figures 1, 3 and Supplementary File 1: Figure S1A were acquired with AxioObserver Z1 epifluorescence microscope equipped with Axiovision software and LD Plan-Neofluar 20x/0.4 Korr Ph 2 M27 objective (Figure 1) or Plan-Apochromat 63x/1.40 oil objective (Figure 3 and Supplementary File 1: Figure S1A). Images for the Figures 4, 5, 6, 7 and Supplementary File 1: Figure S1B and S2 were acquired with LSM 510 Meta confocal microscope equipped with Plan-Apochromat 63x/1.4 oil DIC objective and LSM software.

Image acquisition settings were adjusted so that detected FISH and immunofluorescence signals are above background. To establish the percentage of cells expressing Pab1 and the model RNA presented in Supplementary File 1 Table S2, cells were manually counted and the percentage is the fraction of the cells with FISH or immunofluorescence signal in the total population of cells. The same approach was used to calculate the fraction of cells expressing the model RNA in Pab1-GFP (Figure 4) and Edc3-GFP (Figure 5) strains.

ImageJ was used to quantify P bodies in the Edc3-GFP strain. The images were background subtracted and processed using thresholding, conversion to mask and analyze particles functions.

## 5. Conclusions

We report for the first time accumulation of the mRNA with mammalian codon bias termed the model RNA in yeast mutants, which form constitutive stress granules in the absence of stress. Our data suggest that the accumulation is due to protection of the model RNA from the main degradation path as occurs under stress conditions. At the same time our data do not support the idea that the protection occurs due to the sequestration of the model RNA in stress granules. Moreover, there is no correlation between formation of constitutive stress granules and accumulation of the model RNA up to the fact that in some mutants with efficient stress granule formation the model RNA is downregulated. Thus, our data strongly suggest that the mRNA stabilization and formation of stress granules are driven by different paths which may be activated independently.

## Author Contributions

conceptualization, N.V.K.; experimental design: N.V.K. and A.R.R.; formal analysis, N.V.K.; investigation, N.V.K.; resources, A.R.R. and C.P.; writing—original draft preparation, N.V.K.; writing—review and editing, N.V.K., A.R.R. and C.P.; supervision, A.R.R. and C.P.; funding acquisition, C.P. and A.R.R. All the authors reviewed the results and approved the final version of the manuscript.

## Funding

This research was funded by the program of Région Centre-Val de Loire, “ARD 2020 Biomédicament” (Project A1) and by a grant from la Ligue contre le Cancer comité du Loiret (2014-2015).

## Acknowledgments

We thank everyone who kindly gifted reagents used in this work: Prof. Maria Carmo-Fonseca (Institute of Molecular Medicine, Lisbon, Portugal) for pEGFP-C1 and pGEM-IgM-3MS2, Prof. Ralf-Peter Jansen (Interfaculty Institute of Biochemistry, Tübingen, Germany) for RJP1686 encoding for RFP-ER, Prof. Peter Ponsaerts (University of Antwerp, Antwerp, Belgium) for pGEM4Z/GFP/A64, Dr. Benoit Palancade (Institut Jacques Monod, Paris, France) for *PAB1-GFP* and *EDC3-GFP* strains, Prof. Ambro Van Hoof (The University of Texas Health Science Center at Houston, USA) for *DCP2* mutants, Prof. Maurice S. Swanson (University of Florida, Gainesville, USA) for 1G1 and 4C3 antibodies.

We thank our colleagues from Centre de Biophysique Moléculaire: David Gosset for the help with microscopy, Nadège Hervouet-Coste for introduction to Northern blot, Kevin Moreau for the help with image processing and Christine Mosrin-Huaman for technical advices on yeast genetics. We also thank José Rino and Célia C. Carvalho (Institute of Molecular Medicine, Lisbon, Portugal) for invaluable technical advices on imaging and FISH-IF respectively.

## Conflicts of Interest

The authors declare no conflict of interest. The funders had no role in the design of the study; in the collection, analyses, or interpretation of data; in the writing of the manuscript, or in the decision to publish the results.

## Abbreviations

mRNA: Messenger ribonucleic acid
PKA: Protein kinase A
HPLC: High performance liquid chromatography
FPLC: Fast protein liquid chromatography
(y)(E)GFP: (yeast) (Enhanced) green fluorescent protein
ORF: Open reading frame
UTR: Untranslated region
CSC: Codon stabilization coefficient
NTF: Non-transformed
FISH: Fluorescent *in situ* hybridization
IF: Immunofluorescence
ER: Endoplasmic reticulum
RFP: Red fluorescent protein
PCR: Polymerase chain reaction
DNA: Deoxyribonucleic acid
FCC: Frozen competent cells
PBS: Phosphate-buffered saline
VRC: Ribonucleoside vanadyl complex
LSM: Laser scanning microscope

