## Supplementary File 1 for "mRNA with mammalian codon bias accumulates in yeast mutants with constitutive stress granules"

#### Supplementary figure legends:

**Figure S1. The model RNA is expressed at different levels within the cell population and accumulates in *rtc2Δ* and *cho2Δ* mutants in a path linked to Dcp2 activity.** The model RNA was expressed from episomal plasmid in Tet-Off system in the yeast strains indicated below. (A and B): The model RNA was detected by FISH in the wild-type strain (BY4741), (A) and in *PAB1-GFP* strain (B) using epifluorescence (A) and confocal (B) microscopy, as described in the materials and methods. Uninduced cells were used to control for unspecific labeling. Scale bars are 5 µm. In panel B, the same group of induced cells was imaged at two acquisition settings adjusted either for imaging of cells with medium to low expression of the model RNA (AS1) or cells with high expression of the model RNA (AS2). Uninduced cells were imaged with the settings AS1. Note that panel (A) aims to simultaneously show several cells with different expression levels of the model RNA so that a specific part of a field is used in which the proportion of the cells with different expression levels is distinct from the average values calculated for this strain and shown in Additional file 1: Table S2. (C) The model RNA was detected by northern blot with the probe to *EGFP* using total RNA from wild type strain (w.t., BY4741) and mutants (indicated above each line). Two independent clones were used for *dcp2-7 rtc2Δ* and *dcp2-7 rtc2Δ cho2Δ* mutants. Ribosomal RNA (25S and 18S) was visualized by ethidium bromide staining and served as endogenous control. RNA from uninduced cells (dox) served as control for unspecific hybridization. The white vertical line indicates that the left and right part of the image are taken from the same gel with the same exposure with the middle part removed. Full-length gel and blot for the cropped images are shown in Additional file 1: Fig. S3B.

**Figure S2. Expression of the model RNA downregulates RFP-ER.** The model RNA was expressed as in Additional file 1: Fig. S1 in the wild type (BY4741, W.T.) and the mutants (indicated on the left) also transformed with the plasmid expressing RFP-ER. The model RNA and RFP-ER were detected by FISH-IF. Uninduced cells (W.T. dox) and BY4741 expressing the model RNA only (W.T. no RFP-ER) were used to control for unspecific labeling. Scale bar is 5 µm.

**Figure S3. Full-length gels and blots for the cropped images.** Full-length gels and blots for the cropped images shown on Fig. 2 (A) and Additional file 1: Fig. S1 (B). Arrows indicate the lines which were taken for the cropped images.

### Figure S1

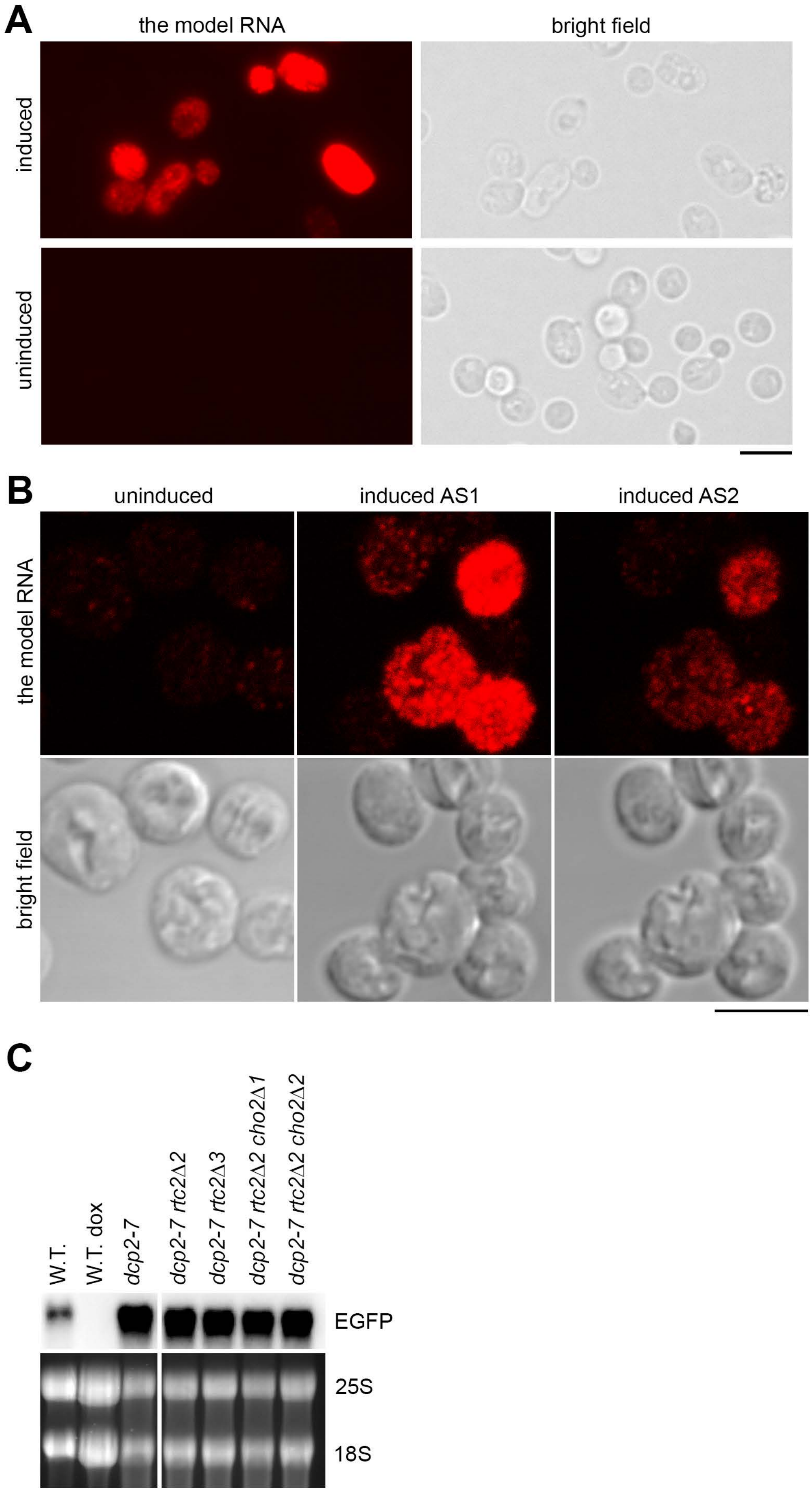
A
the model RNA
bright field
induced
uninduced
B
uninduced
induced AS1
induced AS2
the model RNA
bright field
C
W.T.
W.T. dox
dcp2-7
dcp2-7 rtc2Δ2
dcp2-7 rtc2Δ3
dcp2-7 rtc2Δ2 cho2Δ1
dcp2-7 rtc2Δ2 cho2Δ2
EGFP
25S
18S

### Figure S2

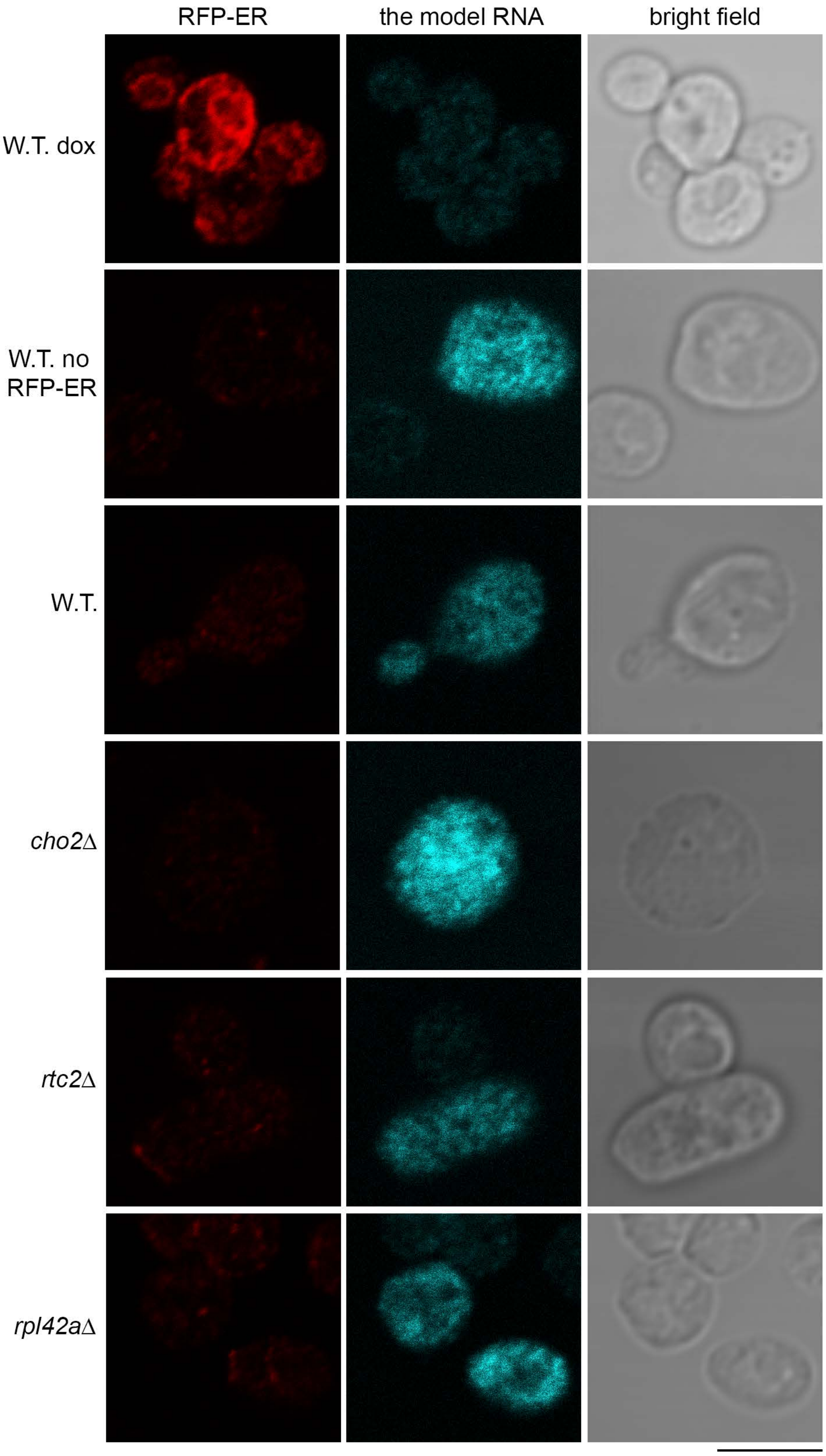
RFP-ER
the model RNA
bright field
W.T. dox
W.T. no RFP-ER
W.T.
*cho2*Δ
*rtc2*Δ
*rpl42a*Δ

Figure S3

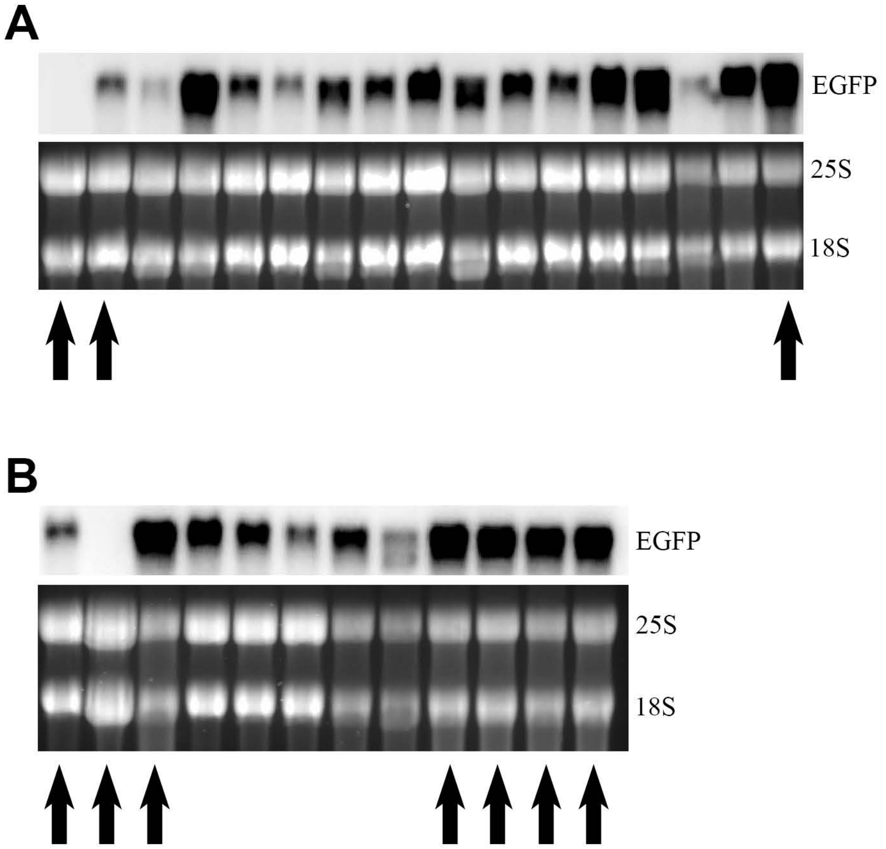

#### Table S1. The model RNA accumulates in mutants with constitutive stress granules

| | Fold increase relative to the w.t.** | | | Consitutive stress granules*** | | Elevated P-bodies*** | | Stress granules and PB*** | |
|---|---|---|---|---|---|---|---|---|---|
| | **Experiment1** | **Experiment 2** | **Experiment 3** | **Cells with SGs (%)** | **Cells with 2+ SGs (%)** | **Av size (uM)** | **Av foci/cell** | **Co-localized** | **Distinct** |
| BY4741 | 1 | 1 | 1 | < 0.2 | 0 | 0.09 | 0.40 | | |
| *mft1 Δ* | 0.8 | 0.5 | 0.48 | 33.00 | 22.00 | 0.14 | 0.85 | | * |
| ***xrn1 Δ*** | **3.8** | **6.9** | **4.86** | 35.00 | 20.00 | | | * | |
| *atg11 Δ* | 0.8 | 1.7 | 2.61 | 18.00 | 0.00 | | | * | |
| *atg15 Δ* | 0.1 | 0.9 | 1.44 | | | | | | |
| ***rlf2 Δ*** | **2.3** | **2.2** | **3.07** | 39.00 | 16.70 | 0.12 | 1.40 | * | |
| *pgd1 Δ* | 0.1 | 2 | 0.54 | 23.00 | 7.90 | 0.14 | 2.00 | * | |
| *kcs1 Δ* | 0.8 | 3.8 | 1.11 | 12.20 | 10.20 | 0.20 | 1.98 | * | |
| *yil001w Δ* | 0.1 | 2.4 | 3.56 | 28.00 | 0.00 | 0.10 | 0.89 | | |
| *swh1 Δ* | 0.2 | 2.5 | 1.42 | 2.70 | 0.50 | 0.14 | 1.47 | * | |
| *swd3 Δ* | 0.3 | 1.8 | 2.34 | 6.70 | 1.00 | 0.14 | 1.55 | * | |
| ***rtc2 Δ*** | **5.8** | **6.2** | **4.32** | 32.00 | 21.10 | | | * | |
| ***cho2 Δ*** | **3.9** | **6.6** | **2.63** | 31.00 | 17.00 | | | * | |
| *dcp2 Δ* | 2.7 | 0.8 | 1.23 | | | | | | |
| ***dcp2-7*** | **2.6** | **5.2** | **2.08** | | | | | | |
| ***rpl42a Δ*** | NT | **8.3** | **4.06** | 8.60 | 2.90 | 0.14 | 1.68 | * | |
| *vma2 Δ* | 0.4 | NT | 2.11 | 22.00 | 16.00 | | | * | |

Several mutants in BY4741 genetic background were tested for ability to accumulate the model RNA. Where appropriate the phenotype of co-localized or distinct stress granules and P bodies is denoted with the asterisk (*).

** Accumulation of the model RNA was tested by quantitative Northern blot and was expressed as fold increase of its amount in the mutants relative to the wild type. Data showing systematic accumulation of the model RNA are highlighted in bold.

*** Data presented in these columns are reproduced from the Table S3 or are according to the Table S2 from (Buchan et al. 2013).

**Table S2. Accumulation of the model RNA does not lead to uniform expression within cell population.**

| Strain | Model RNA expression | | Pab1 expression | Total number of cells |
|---|---|---|---|---|
| | % total | % high | | |
| W.T. | 54% | 16% | 96% | 577 |
| *dcp2-7* | 50% | 25% | 24% | 103 |
| *cho2* Δ | 78% | 27% | 98% | 263 |
| *rpl42a* Δ | 74% | 40% | 91% | 125 |
| *rtc2* Δ | 70% | 17% | 100% | 141 |

Indicated are percent of cells in total cell population expressing the model RNA (% total), expressing it at the very high level (% high) as assesed by FISH and expressing Pab1 (Pab1 expression) as assesed by immunofluorescence.

#### Table S3. Expression of the model RNA does not affect the size and number of P bodies in Edc3-GFP strain

| Model RNA expression | Average foci size, $\mu m^2$ | SD | Average foci/cell | SD | Total number of cells |
|---|---|---|---|---|---|
| uninduced | 0.064 | 0.011 | 0.97 | 0.15 | 93 |
| induced | 0.086 | 0.029 | 0.95 | 0.49 | 88 |

#### Table S4. Primers used in this study.

| Pair number | Primer sequence Forward/ Reverse | Purpose |
|---|---|---|
| 1 | CGGAATTCTTAATCTCGATATCCGTACACCATCAG/ CGGGATCCAGAGGCGGTACCGTCAGCGATTCCGTACCCTGATGG | cloning |
| 2 | CGCAAGCTTCAGTTCGAGTTTATCATTATC/ GCCCCTGCAGGTTTGTTTGTTTATGTGTG | cloning |
| 3 | CGCGGATCCATCATGTAATTAGTTATG/ GCCGAGCTCCAAATTAAAGCCTTCGAG | cloning |
| 4 | AATGAATCGGCCAACGCGCGGGGAGAGGCGGTTTGCGTATTGGGCGCTCTTCCGCTGATCGTACTTGTTACCCATC/ CGCCGCAGCCGAACGACCGAGCGCAGCGAGTCAGTGAGCGAGGAGATCCAATATCAAAGGAAAT | cloning |
| 5 | AACCCAGACGTAATAGCCTCAAGCGAGCATCCCTAAATTTCTGCCATGCGTACGCTGCAGGTCGAC/ AGTAAGTTTGAGTTTTAAGTGGAAAGATTAATAGAAGCCAACTAGTTAATCGATGAATTCGAGCTCG | *RTC2* gene deletion |
| 6 | TTTCGAGTGATTTTCTTAGTGACAAAGCTTTTTCTTCATCTGTAGATGCGTACGCTGCAGGTCGAC/ TGAATCCTAGTACTTTTTAAATATATATACTCAAAAAAAAAAAAACTCAATCGATGAATTCGAGCTCG | *CHO2* gene deletion |
| 7 | TAGTTTCGAATAAACACACATAAACAAACAAACCTGCAGGAAAAATGTCTAAAGGTGAAGAATT/ ATGGGCTAGCTCGTACCCTGATGGTGTACGGATATCGAGATTATTTGTACAATTCATCCA | cloning |
| 8 | TAGTTTCGAATAAACACACATAAACAAACAAACCTGCAGGCGCCACCATGGTGAGCAAGGGCGAGGAG/ ATGGGCTAGCTCGTACCCTGATGGTGTACGGATATCGAGATTACTTGTACAGCTCGTCCATGC | cloning |
| 9 | TAGTTTCGAATAAACACACATAAACAAACAAACCTGCAGGAAAAATGGTGAGCAAGGGCGAGGAG/ ATGGGCTAGCTCGTACCCTGATGGTGTACGGATATCGAGATTACTTGTACAGCTCGTCCATGC | cloning |
